# Cloning of Human ABCB11 Gene in *E. coli* required the removal of an Intragenic Pribnow-Schaller Box before it’s Insertion into Genomic Safe Harbor AAVS1 Site using CRISPR Cas9

**DOI:** 10.1101/2020.09.05.284125

**Authors:** Nisha Vats, Madhusudana Girija Sanal, Senthil Kumar Venugopal, Pankaj Taneja, Shiv Kumar Sarin

**Affiliations:** Institute of Liver and Biliary Sciences, D1 Vasant Kunj, New Delhi; South Asian University, Akbar Bhawan, Chanakyapuri, New Delhi; Sharda University, Plot No. 32, 34, Knowledge Park III, Greater Noida, UP

**Author notes:** **Corresponding Author:** Madhusudana Girija, Sanal MBBS, PhD, Assistant Professor, Department of Molecular & Cellular Medicine, Institute of Liver and Biliary Sciences, D1 Vasant Kunj, New Delhi-110070.

## Abstract

**Background:** Genomic safe harbors are sites in the genome which are safe for gene insertion such that the inserted gene will function properly, and the disruption of the genomic location doesn’t cause any foreseeable risk to the host. The AAVS1 site is the site which is disrupted upon integration of Adeno Associated Virus (AAV) and is considered a ‘safe-harbor’ in human genome because about one third of humans are infected with AAV and so far there is no apodictic evidence that AAV is pathogenic or disruption of AAVS1 causes any disease in man. Therefore, we chose to target AAVS1 site for the insertion of ABCB11, a bile acid transporter which is defective in Progressive Familial Intra Hepatic Cholestasis Type-2 (PFIC-2), a lethal disease of children where cytotoxic bile salts accumulate inside hepatocytes killing them and eventually the patient.

**Methods:** We used CRISPR Cas9 a genome editing tool to insert ABCB11 gene at AAVS1 site in human cell-lines.

**Results:** We found that human ABCB11 sequence has a “Pribnow- Schaller Box” which allows its expression in bacteria and expression of ABCB11 protein which is toxic to *E. coli* and the removal of the same was required for successful cloning. We inserted ABCB11 at AAVS1 site in HEK 293T using CRISPR-Cas9 tool. We also found that ABCB11 protein has similarity with *E. coli* Endotoxin (Lipid A) Transporter MsbA.

**Conclusion:** We inserted ABCB11 at AAVS1 site using CRISPR-Cas9, however, the frequency of homologous recombination was very low for this approach to be successful in-vivo (Figure: pictorial abstract).

**Pictorial Abstract:** ABCB11 gene (which codes the transporter of human bile salts) is targeted to AAVS1 site using a construct which has 5’ and 3’ overhangs which are homologous to the AAVS1 site. A Pribnow box was detected inside ABCB11 gene which allowed the gene to transcribe in E. Coli causing bacterial lysis probably through competitive replacement of a homologous transporter protein in E. Coli (E. coli Endotoxin (Lipid A) Transporter) MsbA, resulting in Lipid A (L) accumulation inside the bacteria.

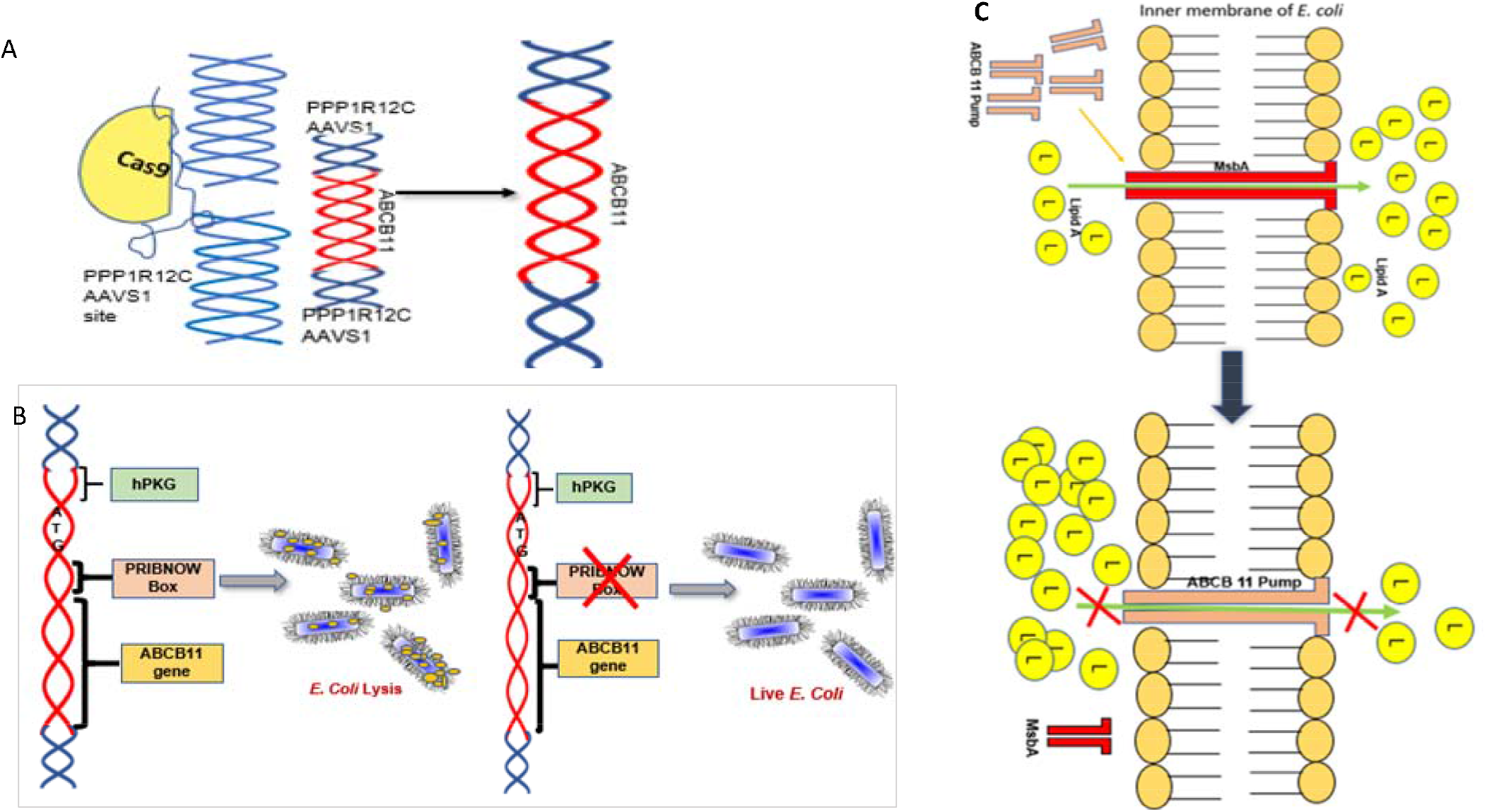

## Introduction

Progressive familial intrahepatic cholestasis type-2 (PFIC2), a severe liver disease which is familial, neonatal, progressive and often fatal which results from a mutation of ATP Binding Cassette Subfamily B Member 11 (*ABCB11*) gene which codes for an ABC transporter Bile Salt Export Pump (BSEP) (1,2). Mutations in *ABCB11* gene results in accumulation of cytotoxic taurocholate and other cholate conjugates leading to progressive hepatocyte destruction (1). Currently the definitive cute for PFIC2 is liver transplantation which is limited by suitable donor organs. Gene therapy, allogenic hepatocyte transplantation (3) and autologous transplantation of hepatocytes/liver organoids differentiated from ‘gene corrected’ induced pluripotent cells could be future options (4,5). Adeno Associated Virus (AAV) is so ubiquitous in man and animals that about 30% of the world population are positive for this virus and to date no disease is proven to be associated with this virus (6–9). In our study we inserted ABCB11 gene at AAVS1 site using CRISPR-Cas9 tool (Figure-1) in HEK293T cells and a fibroblast line.

**Figure-1.**
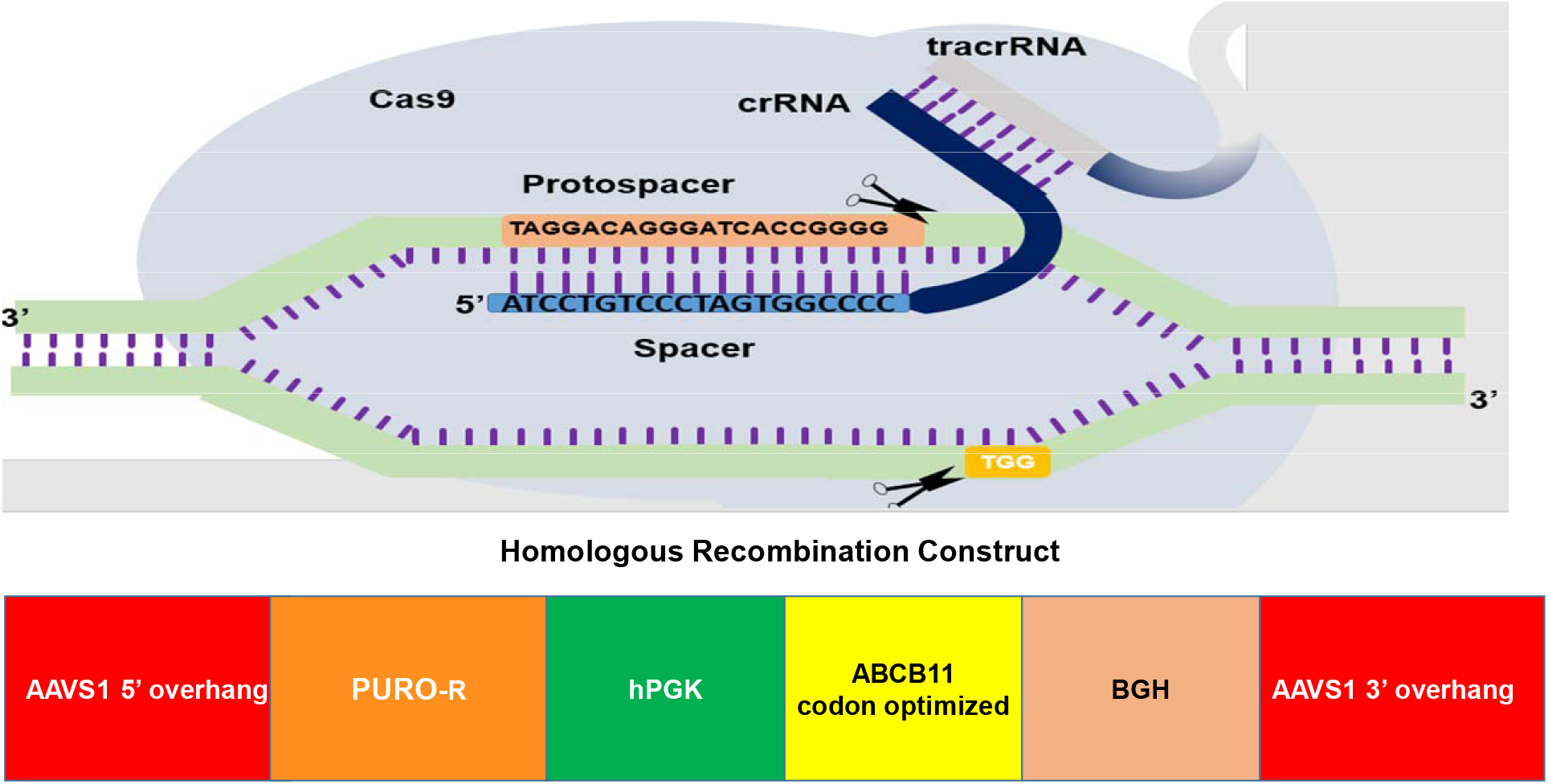
The Cas9-sgRNAs designed and the recombination cassette (donor vector). The gene of interest (ABCB11) was flanked by 5’ and 3’ overhangs which are homologous to the AAVS1 site in chromosome 19. ABCB11 gene was driven by human promoter phospho-glycerol kinase.

## Materials and Methods

### Cloning of ABCB11 Gene

We PCR amplified the 3966 bp ABCB11 (from cDNA prepared from total RNA of human liver tissue) using multiple overlapping primers (Supp.Table-1) which were assembled by overlap extension PCR (10). Phusion DNA polymerase (NEB, US Cat. #M0530L) was used as per the manufacturers protocol. Annealing temperate for all PCRs unless otherwise stated was 60°C. The product was cloned in ‘donor’ vector having AAVS1 recombination overhangs on both ends and ampicillin resistance for selection (Supp. Figure-1). The upstream (5’) overhang sequence (803 bp) in the vector was homologous to a sequence (NCBI Sequence ID: NC_000019.10, 55115768 to 55116570) inside the PPP1R12C gene (Figure-1) which would be in-frame with a 2A ribosome skip sequence and puromycin resistance gene if insertion of the construct happens by homologous recombination with the target site. This is region was followed by a poly adenylation signal and the downstream (3’) overhang sequence which was the continuation of the 5’ overhang (837 bp). We inserted the ABCB11 sequence driven by human phospho-glycerol kinase (hPGK) promoter between these overhang sequences. DH5α cells, JM 109 and One Shot™ Stbl3™ Chemically Competent *E. coli* cells (ThermoFisher Catalog#:C737303) were used for transformation and cloning.

### Bioinformatics

Bacterial Promoter Prediction was done using BPROM-a prediction tool for bacterial promoters (11). DNA or protein sequence comparisons were done using the appropriate tool from NCBI BLAST platform (12). Primers were designed manually or using NCBI Primer Blast (13).

### gRNA

We designed two guide RNAs targeting AAVS1 site (Figure-1) and cloned in two vectors expressing SPCas9 and SACas9 at BsaI sites (Supp. Figure-2) following the standard gRNA design, cloning protocols and resources (14). Off-target analysis was done using the tool Custom Alt-R^®^ CRISPR-Cas9 guide RNA(15).

### Western Blot

Whole cell extracts were run on 10% SDS-PAGE and transferred to a polyvinylidene difluoride membrane using a transfer apparatus following the standard protocols (Bio-Rad). After incubation with 5% nonfat milk in TBST (10 mM Tris, pH 8.0, 150 mM NaCl, 0.5% Tween 20) for 60 min, the membrane was washed once with TBST and incubated with antibodies against human ABCB11 (Affinity, Catalog #DF 9278) 1: 2000 dilution; human β-actin (Santa Cruz Cat.# SC4778), dilution 1:1000. The membrane was washed and incubated with a 1:5000 dilution of horseradish peroxidase-conjugated anti-rabbit (Santa Cruz Cat# SC-2004)/anti-mouse antibodies (Cat.#SC-2005) for 2 h. Blots were washed with TBST four times and developed with the ECL system (Bio-Rad, US Cat.#170-5060) according to the manufacturer’s protocols.

### Cell Culture

HEK293T, HepG2, FS1 (fibroblast) cells were grown in high glucose DMEM (Hi-Media Lab, Mumbai, Cat.# AL111-500ML) supplemented with 10% fetal bovine serum (CellClone, Genetix Biotech Asia, New Delhi, Cat.# CCS-500-SA-U), penicillin and streptomycin (Hi-Media, Mumbai Cat. # A018-5X100ML). When 80% confluent, the cell lines were transfected with Cas9-sgRNA vectors (without the donor vector). 48h post-transfection, about 10000 of these cells were used for comet assay (16) and genomic DNA (gDNA) isolated from the remaining cells was used for T7-endonuclease assay (17) to evaluate the in-vitro ‘DNA cutting’ activity of Cas9-sgRNA construct. Subsequently, we transfected these cells with the donor vector, Cas9-sgRNA vector and a control vector (pEGFPN1) in the ratio: 2:1:1 using PEI (18) from Sigma Aldrich, Inc. (CAS #9002-98-6). After 48h the cells were imaged and used for downstream applications. Half of the transfected dishes were serially passaged without puromycin selection for two weeks and DNA and protein were isolated. On the remaining dishes puromycin selection was started following the manufacturer’s protocol (19) after 36h of transfection and puromycin resistant colonies at 8 μg/mL were further cultured in puromycin containing media and gDNA was isolated.

### T7 Endo Nuclease Assay

T7 endo I assay detects heteroduplex DNA that results from annealing DNA stands that have been modified after a sgRNA/Cas9 mediated cut to DNA strands without modifications. T7 Endonuclease-1 was purchased from NEB, US (Cat. #NEB #E3321) and was used to digest the PCR products amplified from gDNA extracted from Cas9-sgRNA transfected (test) and un-transfected cells (control) using primers flanking the expected Cas9-sgRNA cut sites following the manufactures protocol (17).

### Comet Assay

Fifty to hundred treated cells were embedded in 0.7% low melting agarose and mounted on a precoated slide and was immersed in alkaline 0.1% SDS solution overnight, neutralized and electrophoresis was done in an alkaline buffer (pH 10) at 0.74 V/ cm for 30 minutes (16).

## Results

### Human ABCB11 Gene/Product is Toxic to *E. coli* Cells

Few ampicillin resistant DH5α *E. coli* colonies which we got after transformation were screened for the insert by colony PCR. One colony was positive for all the fragments of the ABCB11 gene. Sequencing revealed that mutations in ABCB11 (Figure-2a,b). Repeated attempts failed and we considered the possibility of unstable DNA sequences. Therefore, we tried JM109 which gave one positive colony and plasmid was isolated. However, after we soon found the bacteria failing to grow or losing the plasmid on subsequent cultures. Therefore, we transformed One Shot™ Stbl3™ Chemically Competent *E. coli* cells, which are suitable for cloning unstable DNA segments. We got many positive colonies, however, upon overnight culture the bacteria formed a big pellet (partly lysed bacteria) which cannot be resuspended in phosphate buffered saline. Therefore, we concluded that ABCB11 gene/gene product is toxic to bacterial cells. It is possible the ABCB11 being a membrane transporter may be toxic to bacteria.

**Figure-2 a, b.**
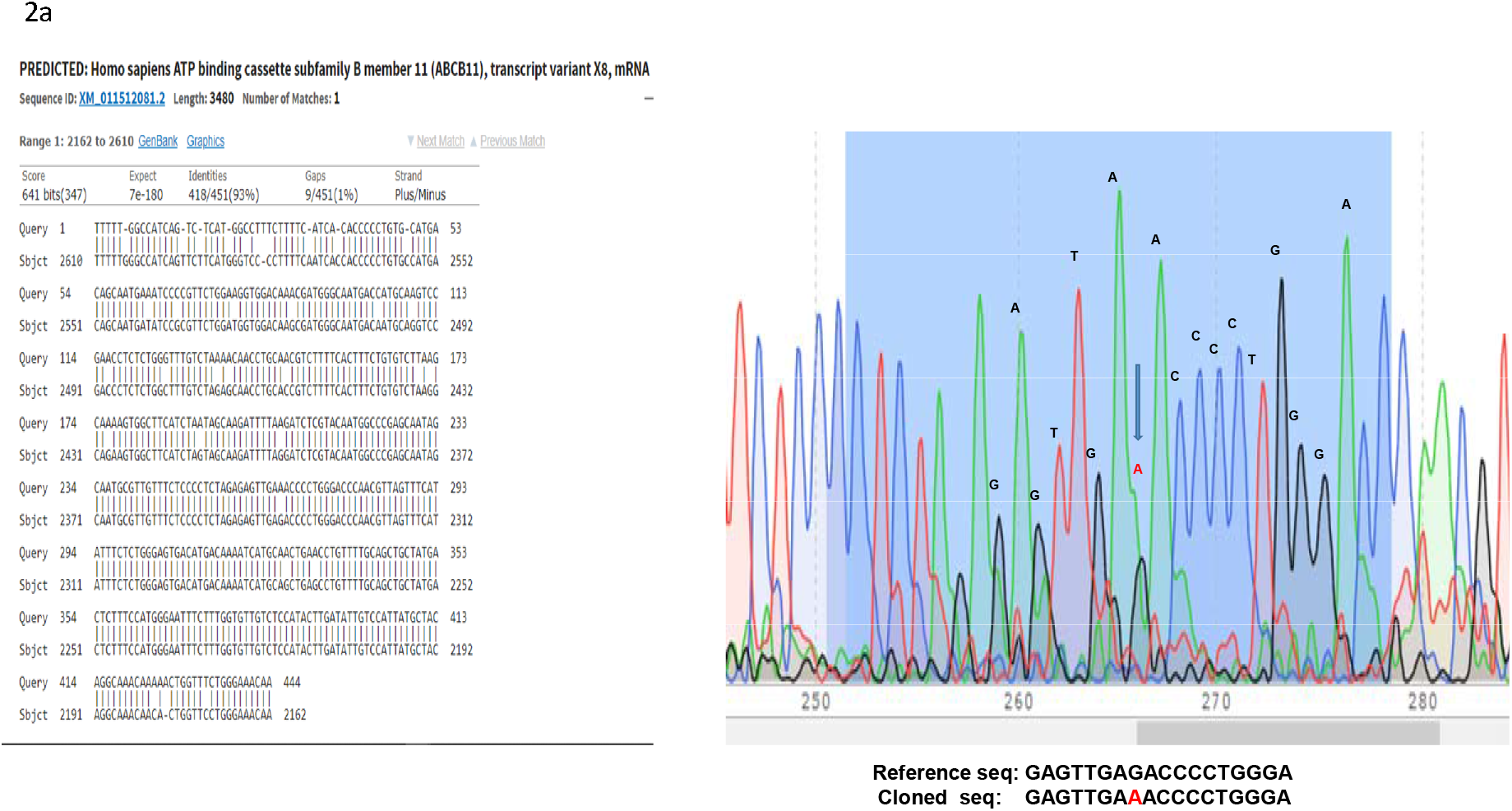
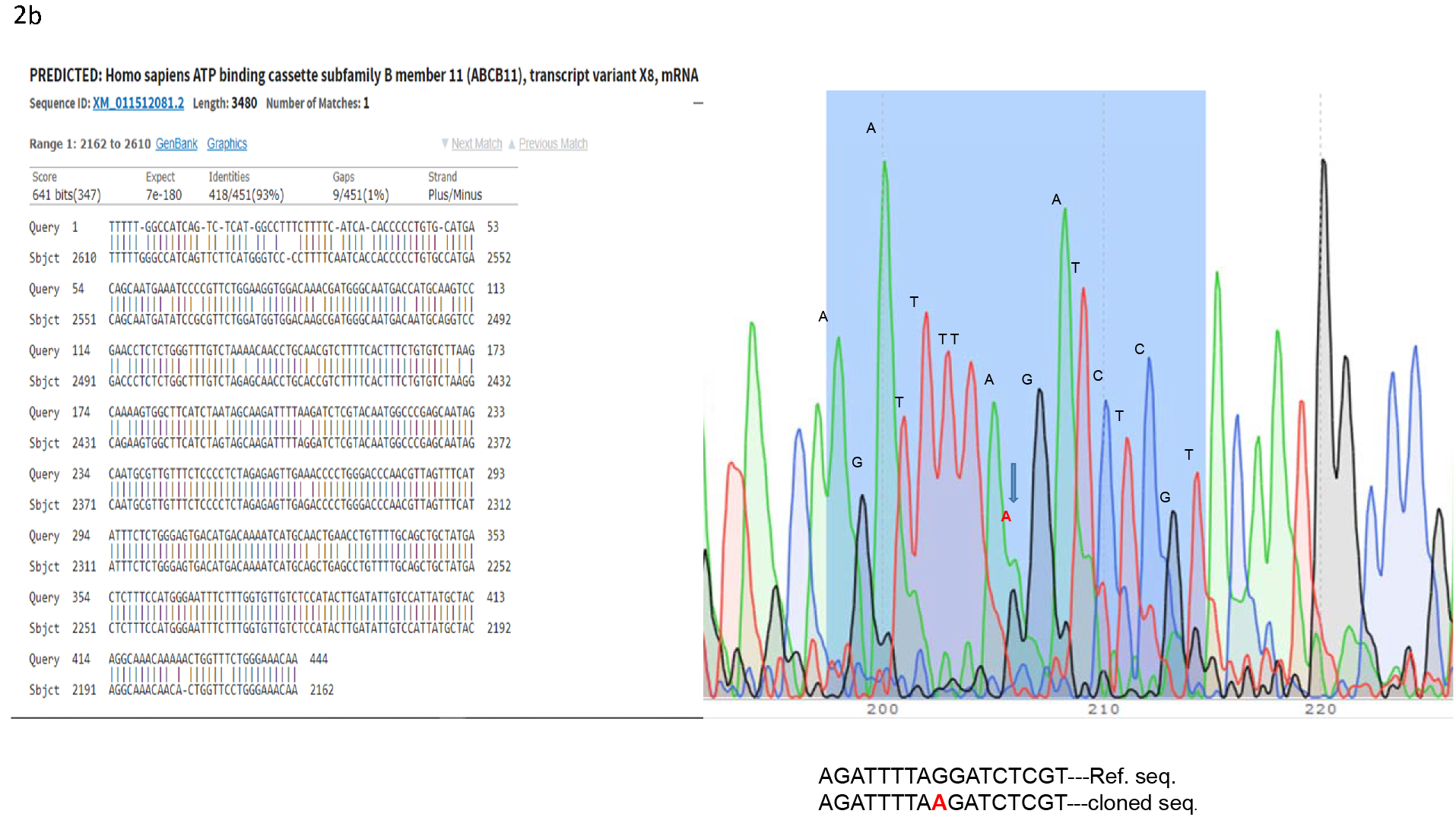
The ABCB gene cloned into *E. Coli* showed mutations despite multiple cloning attempts.

### Identification and Removal of a Pribnow-Schaller Box in human ABCB11 for cloning in *E. coli*

We did a PAGE followed by Coomassie staining to see differential protein expression between ABCB11 donor vector transformed bacteria versus untransformed bacteria (Figure-3a). We found differential expression of a few proteins. We subsequently performed a Western Blot and interestingly antibody against human ABCB11 identified a specific protein over expressed in the transformed cells (Figure-4b). However, in our construct ABCB11 gene was under a eukaryotic promoter. Considering the possibility of some DNA elements which have similarity to bacterial promoters inside the ABCB11 sequence we performed a bioinformatic analysis using BPROM to predict hidden bacterial promoters (Supp.Table-2). The promoter-site (Pribnow-Schaller box tca**tataat**) containing sequence (ggttttgagtcagataaatca**tataat**aat) which we identified was modified to (ggtTTCGAAtcagataaatcaTACAACaat) by PCR using an oligonucleotide primer sequence incorporating the modified sequence and subsequent overlap extension PCR amplification of the entire gene fragment. With this modification we were able to clone ABCB11 coding sequence which was not toxic to bacteria.

**Figure-3.**
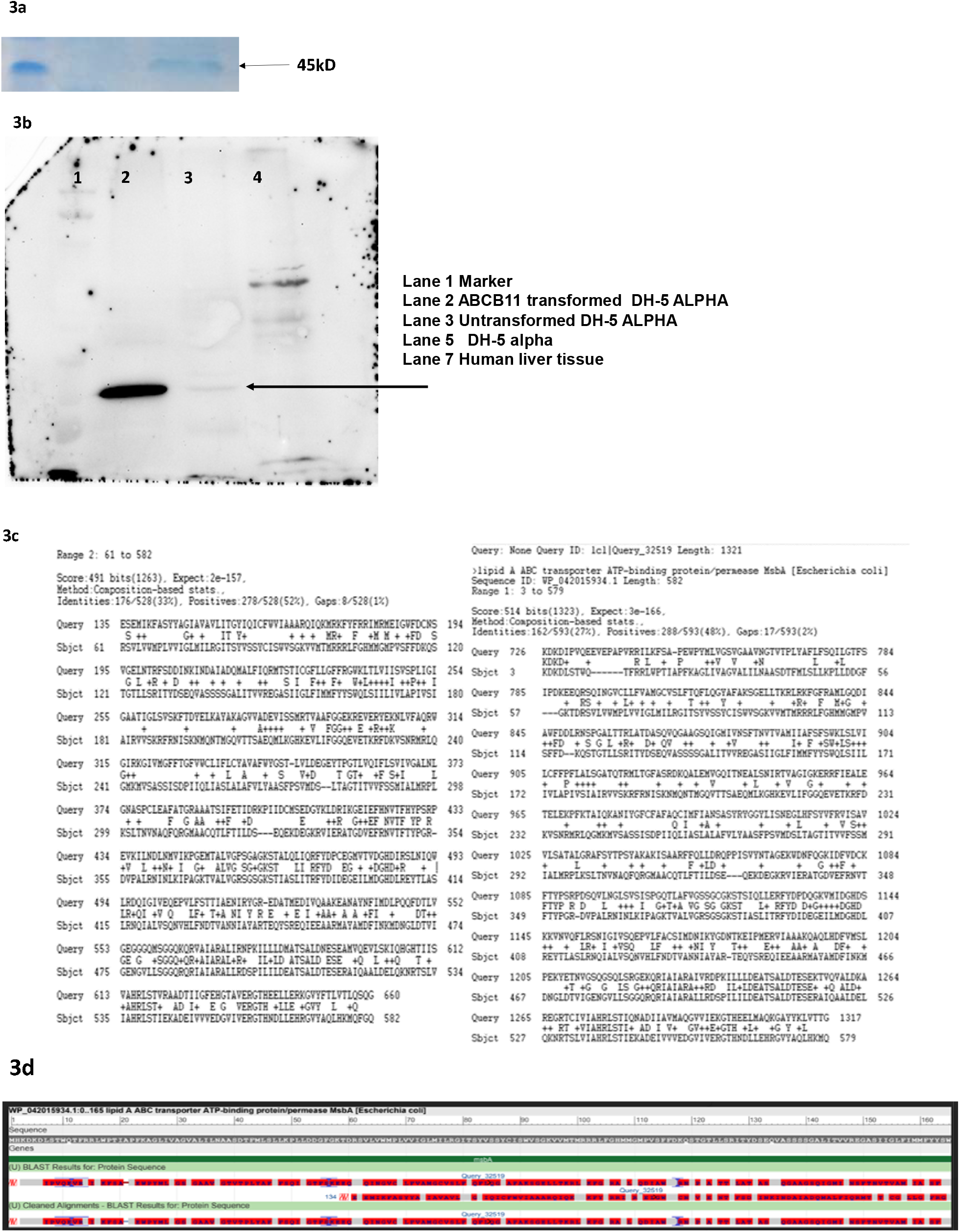
3a: Total protein extract from E. Coli transformed the donor vector on Poly Acryl amide Gel Electrophoresis followed by Coomassie blue staining showed differential expression multiple proteins. 3b: Western Blot with anti-human ABCB11 antibody shows multiple bands including one probably corresponding to MsbA-a bacterial Lipid A transporter. 3c: A bioinformatic analysis (Protein BLAST) revealed ABCB11 and MsbA are sharing conserved domains. 3d: Sequence alignment of ABCB11 and MsbA showing conserved domains

**Figure-4.**
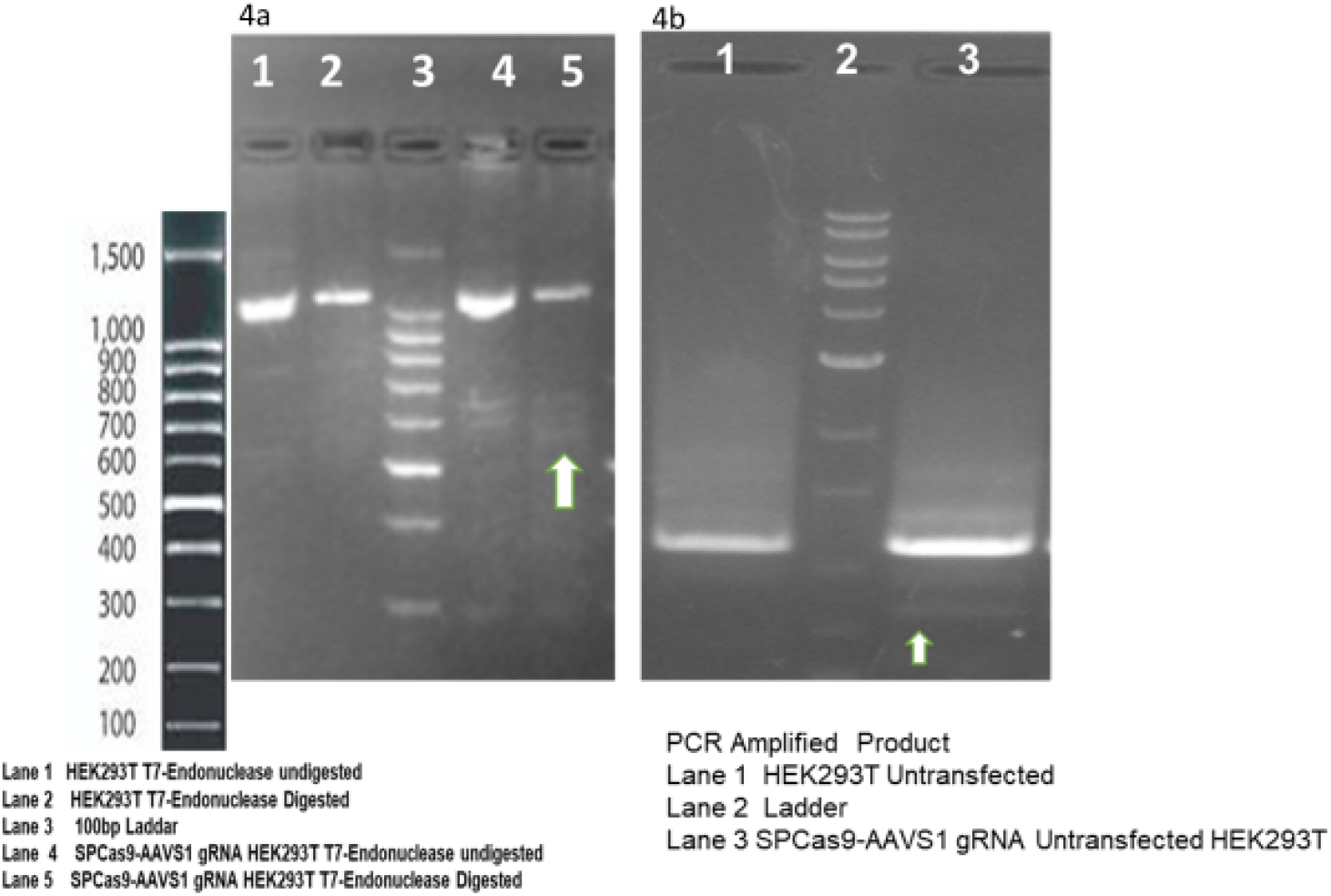
4a: Agarose gel electrophoresis of PCR amplified product after digestion with T7 endonuclease shows bands resulting from the digestion of heteroduplexes 4b: The PCR product without T7 digestion also shows a distinct band resulting from the formation of heteroduplex.

### ABCB11 is similar to *E. coli* Lipid A (Endotoxin) Transporter MsbA

Protein BLAST search identified MsbA (UniProtKB - P60752), a member of the ABC transporter superfamily. MsbA, a 64.46 kD protein, has an important role in *E. coli* Lipid A (endotoxin or LPS) transport (Figure-3c, d). This protein flips core endotoxin from its site of synthesis on the inner leaflet of the inner membrane to the outer leaflet of the inner membrane. Western Blot showed identified a unique band in the donor vector transformed *E. coli* while the untransformed *E. coli* also showed a faint band but specific band at the same position (Fig-3b).

#### Verification of the gDNA cutting activity of Cas9-sgRNA construct

T7 Endonuclease Assay and Comet Assay was used to evaluate the gDNA cutting activity of Cas9-sgRNA. We observed digestion of heteroduplexes at the CRISPR-Cas9 cut sites which were sensitive to T7 endonuclease (Figure-4a). These heteroduplexes were observed on the agarose gel electrophoresis of PCR products as well (Figure-4b). The Cas9-sgRNA damaged the genome of the transfected cells leading to the formation of comet shaped nuclear material upon electrophoresis (Figure-5).

**Figure-5.**
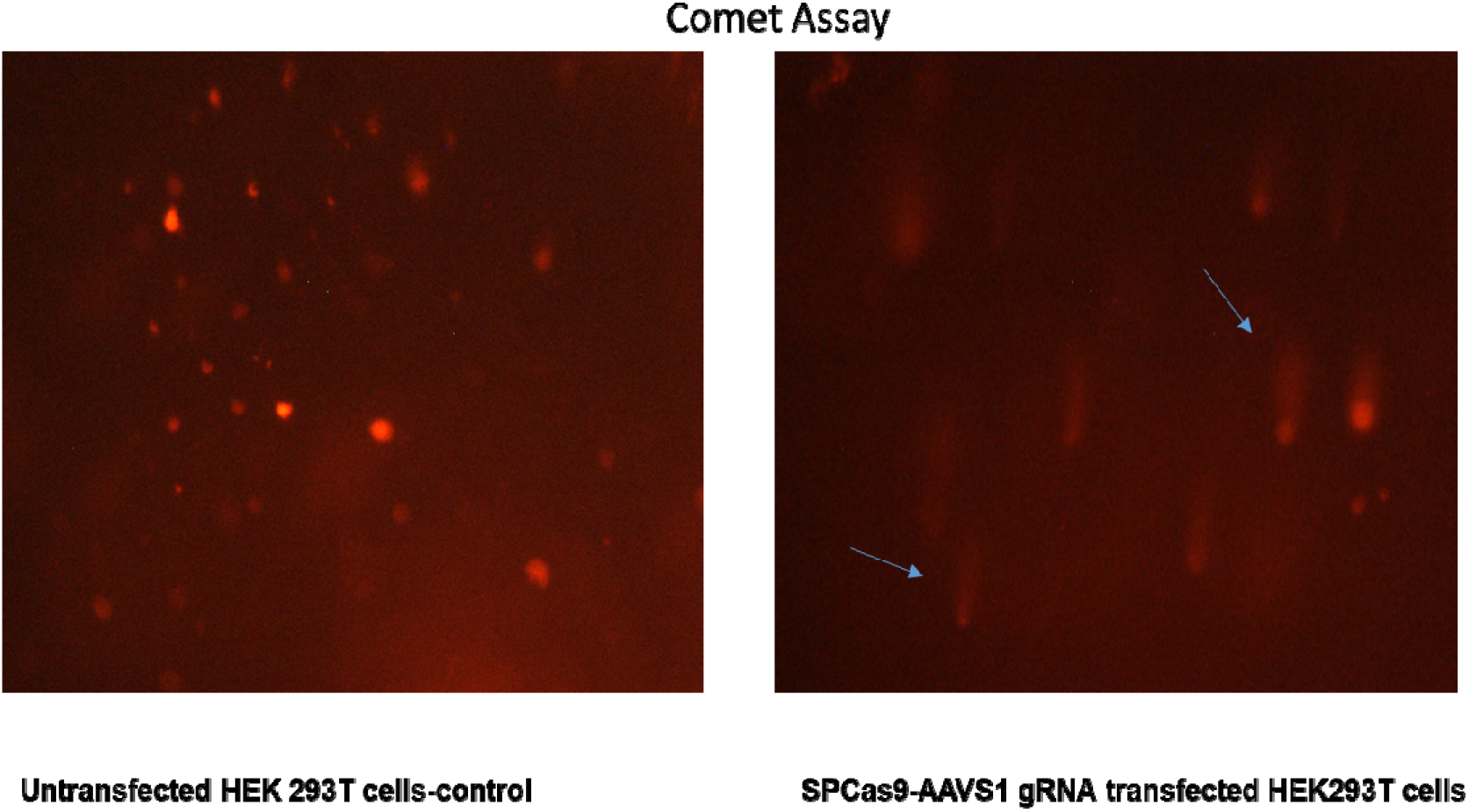
Cas9-sgRNA cause DNA damage which is revealed by a Comet assay. The untreated cells have intact round/oval nuclei while the Cas9-sgRNA treated cells shows a comet shaped nucleus because of DNA damage.

#### Off-Target Analysis

Oligonucleotide PCR primers were designed for bioinformatically predicted off-targets (Supp. Tables-3a, b). We amplified these segments using genomic DNA extracted from the treated cells (48h post-transfection with SPCas9-sgRNA vector) as template. The PCR products were sequenced and analyzed for sequence disruption (Table-1).

**Table-1.**
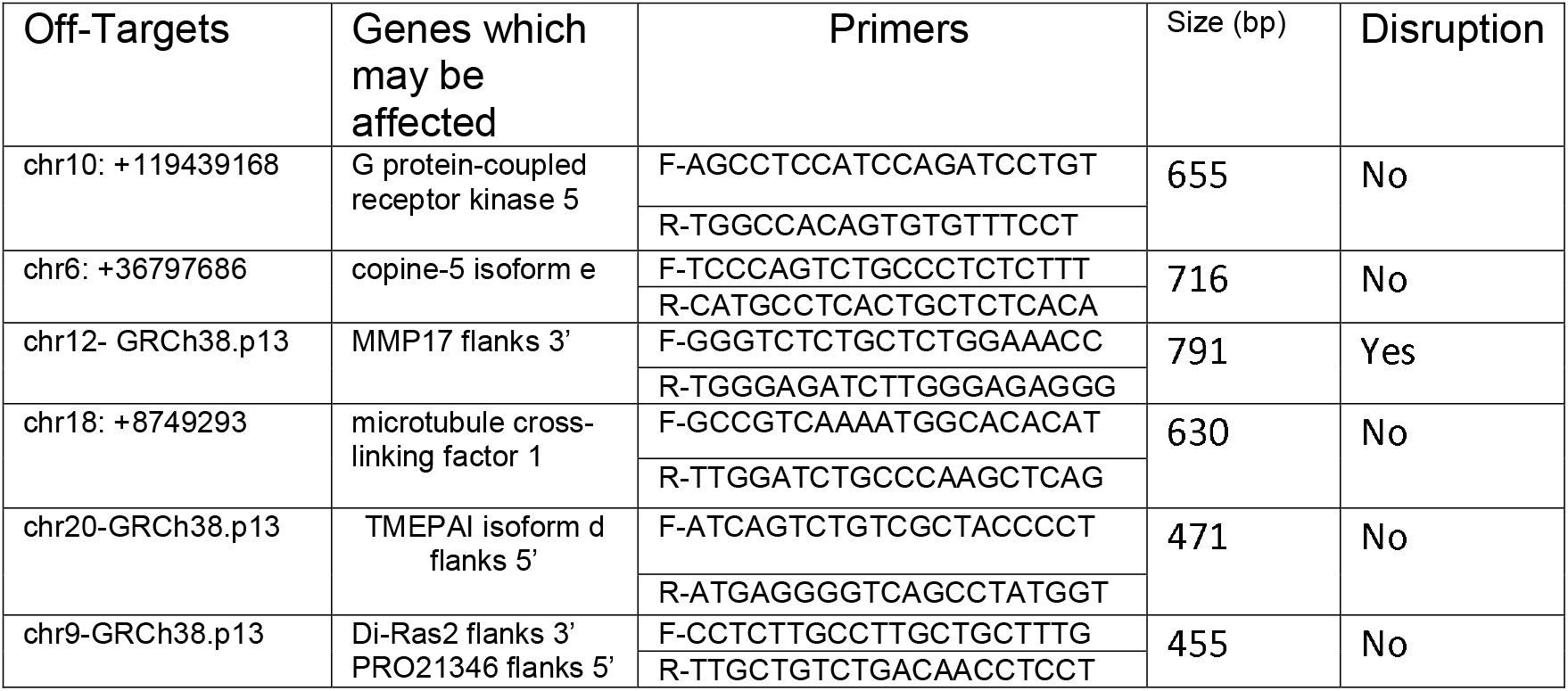
Selected off targets for crRNA GGGGCCACTAGGGACAGGAT (PAM TGG), Oligonucleotide primers designed to amplify these off-target sites to verify off target disruptions

#### Validation of ABCB11 Expression in a Fibroblast Cell Line

The Western Blot done with total protein extract of fibroblasts 48h post-transfection with the ABCB11 donor vector showed the expression of ABCB11 protein (Figure-6a). A fibroblast line was used in these experiments because they don’t express ABCB11 naturally while cell lines such as HEK293T and liver cell lines such as HepG2 express ABCB11.

**Figure-6.**
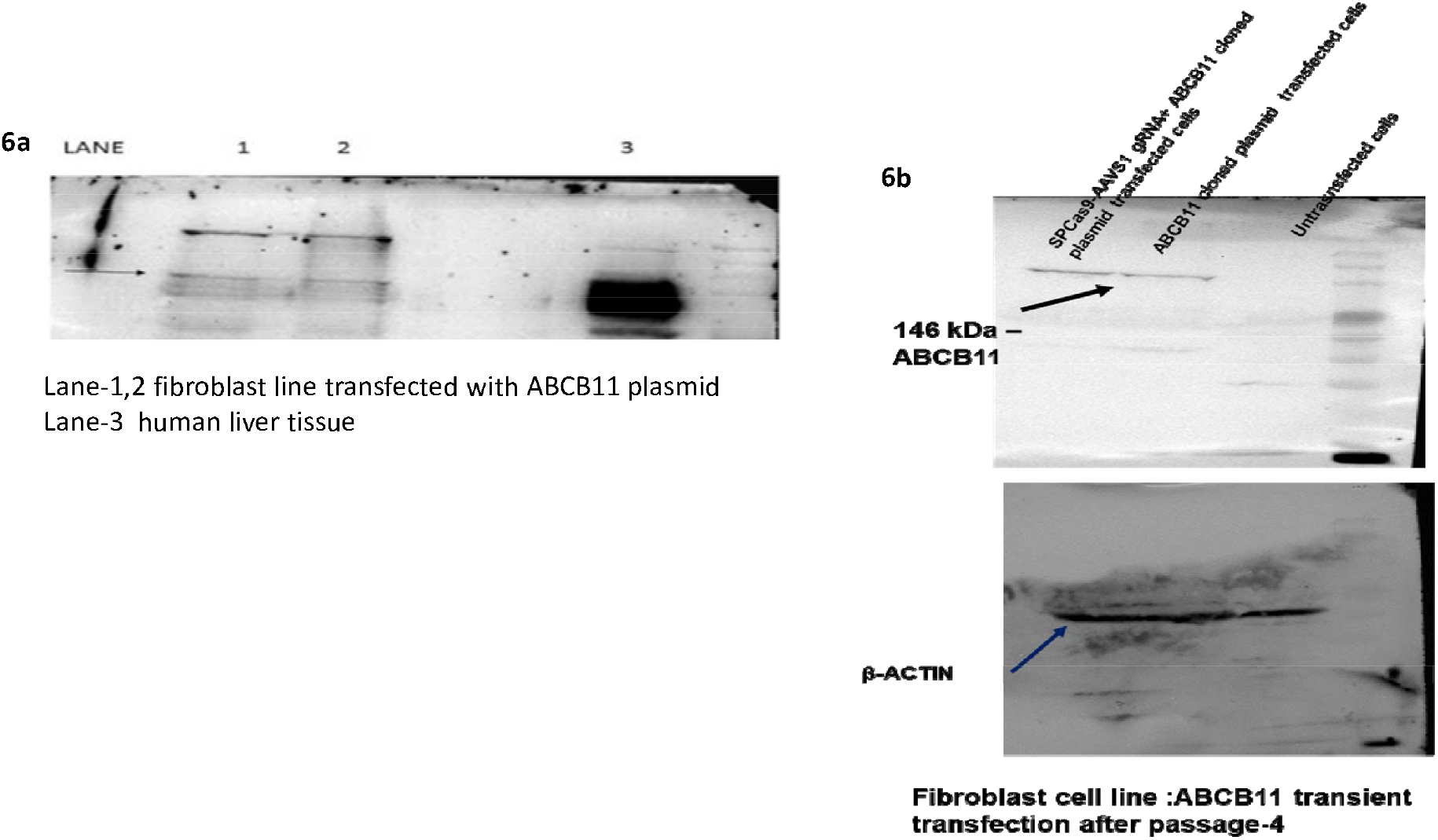
a: Western Blot with anti-human ABCB11 antibody confirmed the expression after transient transfection with the donor vector having ABCB11. b: Western Blot was done using total protein isolate from fibroblasts (which does not naturally express ABCB11) after four passages post-co-transfection of the donor vector with ABCB11 gene and the CRISPR-Cas9-sgRNA vector.

**Figure-7.**
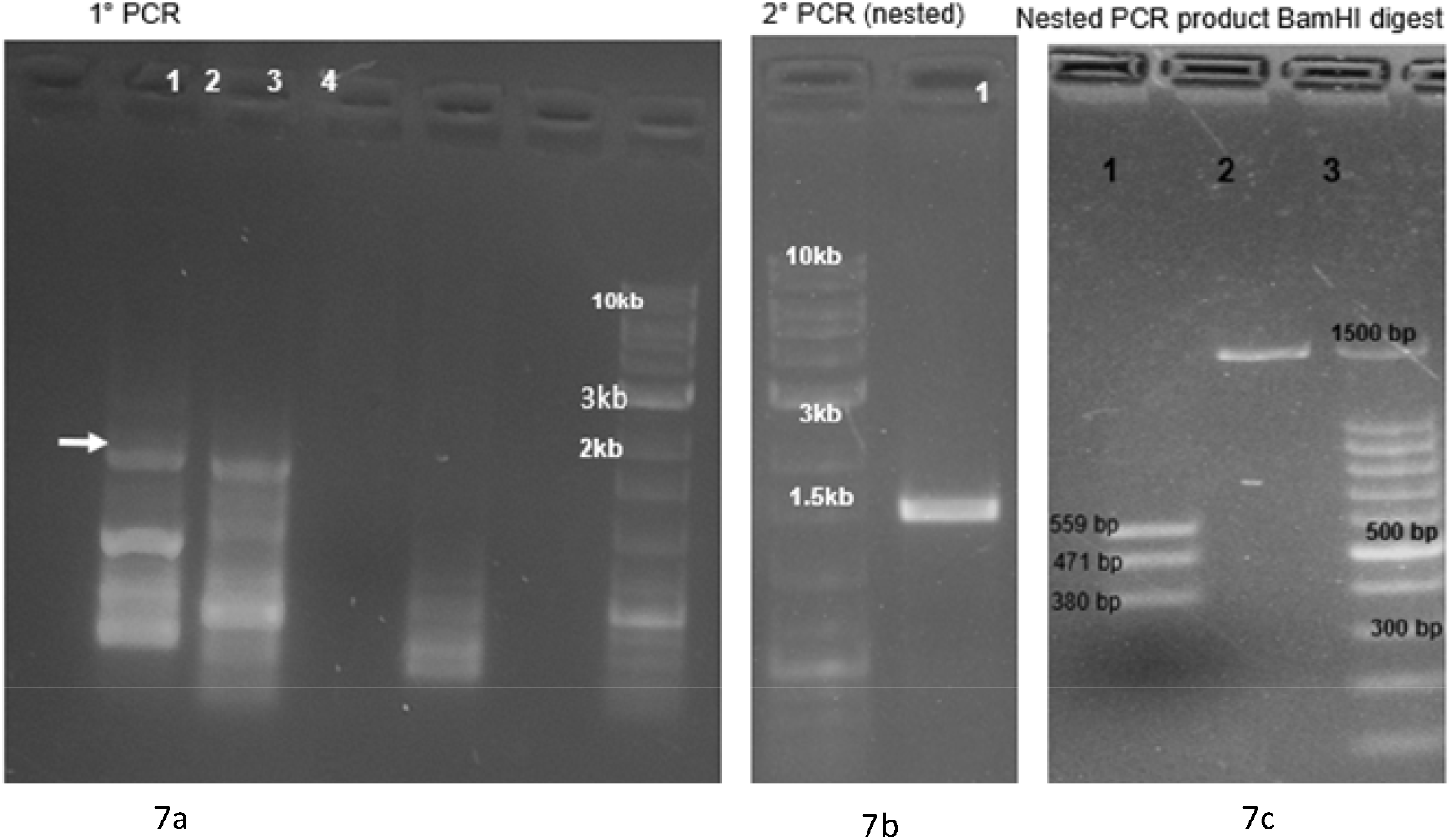
7a: The gDNA isolated from transfected cells (Cas9-sgRNA plasmid alone) after 72h, (Cas9-sgRNA plasmid plus donor vector) after 21 days of puromycin selection were used as PCR templates with a forward primer complementary to a region upstream of the 5’ recombination overhang of the vector and a reverse primer complementary to a sequence in the puromycin resistance gene to amplify a segment spanning from a site in the host cell genomic DNA slightly upstream of the genomic integration site to a segment donated by the donor vector (puromycin resistance gene). This PCR product was used as a template for a nested PCR and product was confirmed by restriction enzyme digestion (BamH1) and sequencing (Primers: Supp. Table-4, Figure-7 b, c).

Western Blot was repeated with total protein extract from fibroblasts after 2 weeks (fourth passage) post-transfection with SPCas9-sgRNA vector together with the donor vector containing ABCB11. This blot also showed the expression of ABCB11 protein (Figure-6b) suggesting the integration of the ABCB11 into the host genome.

#### PCR Amplification and Sequencing to Verify the Integration of ABCB11 to AAVS1 Site

We got only three puromycin resistant colonies upon transfecting about 20 million HEK cells with a transfection efficiency of 70 to 80% in four 6 cm dishes. The gDNA isolated from transfected cells (Cas9-sgRNA plasmid alone) after 72h, (Cas9-sgRNA plasmid plus donor vector) after 21 days of puromycin selection were used as PCR templates with a forward primer complementary to a region upstream of the 5’ recombination overhang of the vector and a reverse primer complementary to a sequence in the puromycin resistance gene to amplify a segment spanning from a site in the host cell genomic DNA slightly upstream of the genomic integration site to a segment donated by the donor vector (puromycin resistance gene). This PCR product was used as a template for a nested PCR and product was confirmed by restriction enzyme digestion (BamH1) and sequencing (Primers: Supp. Table-4, Figure-8a-c). We also PCR amplified parts of ABCB11 using primers (Supp. Table-1) which would give specific PCR products from the inserted cassette, to make sure the gene is not deleted from the cassette integrated to the host cell. PCR products showed the expected sizes confirming that the amplified products are from the cassette and not from the native ABCB11 gene present in the cell line (Figure-8).

**Figure-8.**
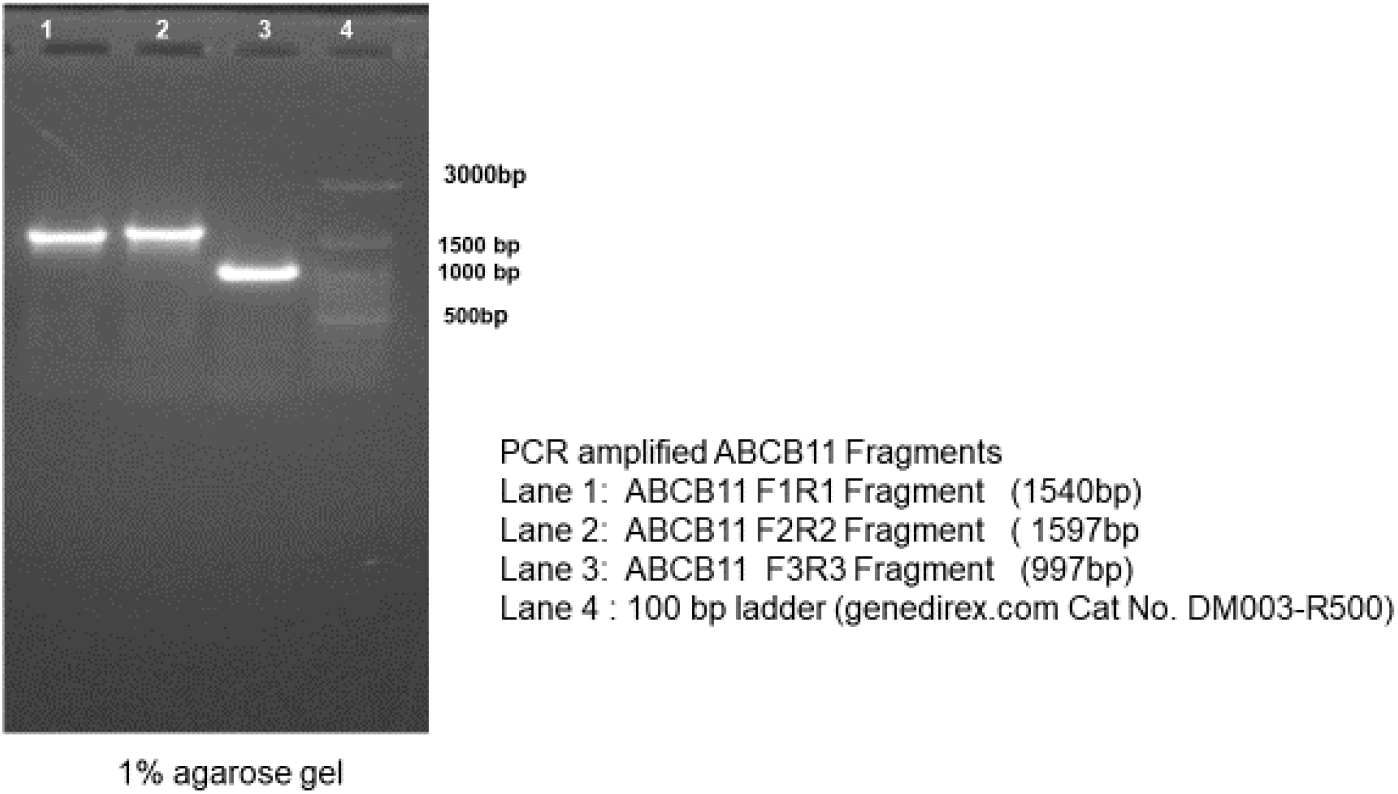
PCR products amplified from the genomic DNA. ABCB11 specific primers were used which would give specific PCR products from the inserted cassette. This was done to make sure the ABCB11 gene was not deleted from the cassette integrated to the host cell genome. The PCR products showed the expected sizes confirming that the amplified products originated from the integrated cassette and not from the native ABCB11 gene present in the cell line.

## Discussion

In our knowledge, this is the first time ABCB11 gene was inserted to AAVS1 safe-harbor using CRISPR-Cas9 technology in human cells. Liver directed gene therapy is another approach and successful in rodents (20). Adeno associated vectors do not integrate and therefore the effect of gene therapy many not last in human beings, especially in infants, as the viral vector dilutes out as the cells proliferate in a growing liver (21). Another approach is transplantation hepatocytes differentiated from gene corrected patient iPSC (4,22,23). AAVS1 site is considered as a ‘safe haven’ in human genome (24) where we chose to insert the gene mutated in PFIC. We could not find any other study which attempted to place ABCB11 gene at AAVS1 site. AAV is a common virus and it is considered non-pathogenic because seroprevalence of wild-type AAV in humans ranges from 40% for AAV8 to 70% for AAV1 and AAV2 and yet we are not aware of any disease caused by AAV (6,7,25). AAV integration into AAVS1 site causes disruption of PPP1R12C (Protein Phosphatase 1 Regulatory Subunit 12C). However, this gene is not associated with any disease (25). A puromycin gene was placed in the donor cassette such that the puromycin gene will be transcribed only if homologous recombination happens. We got only a few puromycin resistant colonies suggesting that homologous recombination was a rare event (^~^10^−7^). This suggests that in-vivo gene therapy using CRISPR-Cas9 technology making use of homology directed gene repair could be difficult. “Targeted Integration and high transgene expression in AAVS1 Transgenic mice after in-vivo hematopoietic stem cell transduction with HDAd5/35++ Vectors”(26) is reported however, to achieve this they used an adenoviral gene delivery system with AAV5 ITRs and AAV35 helper. Integration of AAV/ Cas9 into Cas9 mediated cut sites is a potentially hazardous consequence of this approach (27, 28).

We found that human ABCB11 donor vector transformed bacteria either died or the ABCB11 gene sequence got mutated meaning either the DNA sequence or the ABCB11 protein has some untoward effects on bacteria. We performed a Western Blot and found that in ABCB11 transformed bacterial clones giving a band around 45 kD and HepG2 cells/liver tissue is giving a band at around 140 kD which corresponds to ABCB11. It is interesting to note that untransformed *E. coli* cells are also showing a band although very faint around 45kD suggesting the possibility of a bacterial protein which might have structural similarity to human ABCB11. We performed a bioinformatic search and identified MsbA an *E. coli* protein which functions as a lipid transporter (^~^64kD). MsbA is involved in the transport of bacterial endotoxin-a function like the ABCB11 which transports the bile salts which are lipid derivatives (29,30). Another interesting observation was the identification of a bacterial promoter sequence (Pribnow-Schaller Box) in human ABCB11 causing unexpected expression of ABCB11 protein and bacterial toxicity. This was an important lesson for us because we spent a lot of time trying to clone ABCB11. It is therefore important to search for and eliminate if any bacterial promoter sequences or similar elements are identified for successful cloning of eukaryotic/toxic genes in *E*. coli. Exactly we do not know how ABCB11 caused bacterial toxicity. Possibly expression of ABCB11 in *E. coli* might be destabilizing *E. coli* membrane. ABCB11 being a lipid transporter is a membrane spanning protein.

Alternatively, the human protein, which is similar to the *E. coli* protein might have caused a competition between bacterial transporter MsbA and human protein for membrane incorporation resulting in the accumulation of endotoxin within *E. coli* cells because unlike the bacterial transporter the membrane incorporated ABCB11 might not be able to transport endotoxin out. This raises the possibility that endotoxin is toxic to *E. coli* itself if it is not exported and MsbA can be considered as a drug target which required further validation. The advantage of such an antibiotic is that it will be selective to endotoxin producing microbes. More research is required in this direction. It may be noted that endotoxin producing microbes play an important role in sepsis (31) and diseases such as non-cirrhotic portal hypertension (32–34).

To conclude, inserted ABCB11 at AAVS1 site using CRISPR-Cas9, however, the frequency of homologous recombination was very low as evident from the number of puromycin resistant colonies. With this low efficiency, with the current technology it is unlikely that this approach would be successful in-vivo gene editing.

## Disclosures

### Ethical approval and Informed consent

No human patient or animal material/specimens were used in this work which requires an Institutional Ethics Committee/Institutional Review Board approval to the best of our knowledge.

## Acknowledgement

The corresponding author (Sanal MG) acknowledges the Science and Engineering Research Board (grant # ECR/2015/000275) and Department of Biotechnology, Ministry of Science and Technology, India (grant # BT/PR15116/MED/31/334/2016) Government of India for a limited financial support for about 18 months. The corresponding author thanks Mr. Rahul Saha for his assistance in Western Blot.

## Original data

Original unmodified data of experiments including repeats and failure are uploaded to ‘Dataverse’ (https://dataverse.harvard.edu/).

year = {2020},

version = {DRAFT VERSION},

doi = {10.7910/DVN/CJAODG},

url = {https://doi.org/10.7910/DVN/CJAODG}

