## Supplementary material for "Cloning of Human ABCB11 Gene in *E. coli* required the removal of an Intragenic Pribnow-Schaller Box before it’s Insertion into Genomic Safe Harbor AAVS1 Site using CRISPR Cas9": Supp. figures

### Slide 1
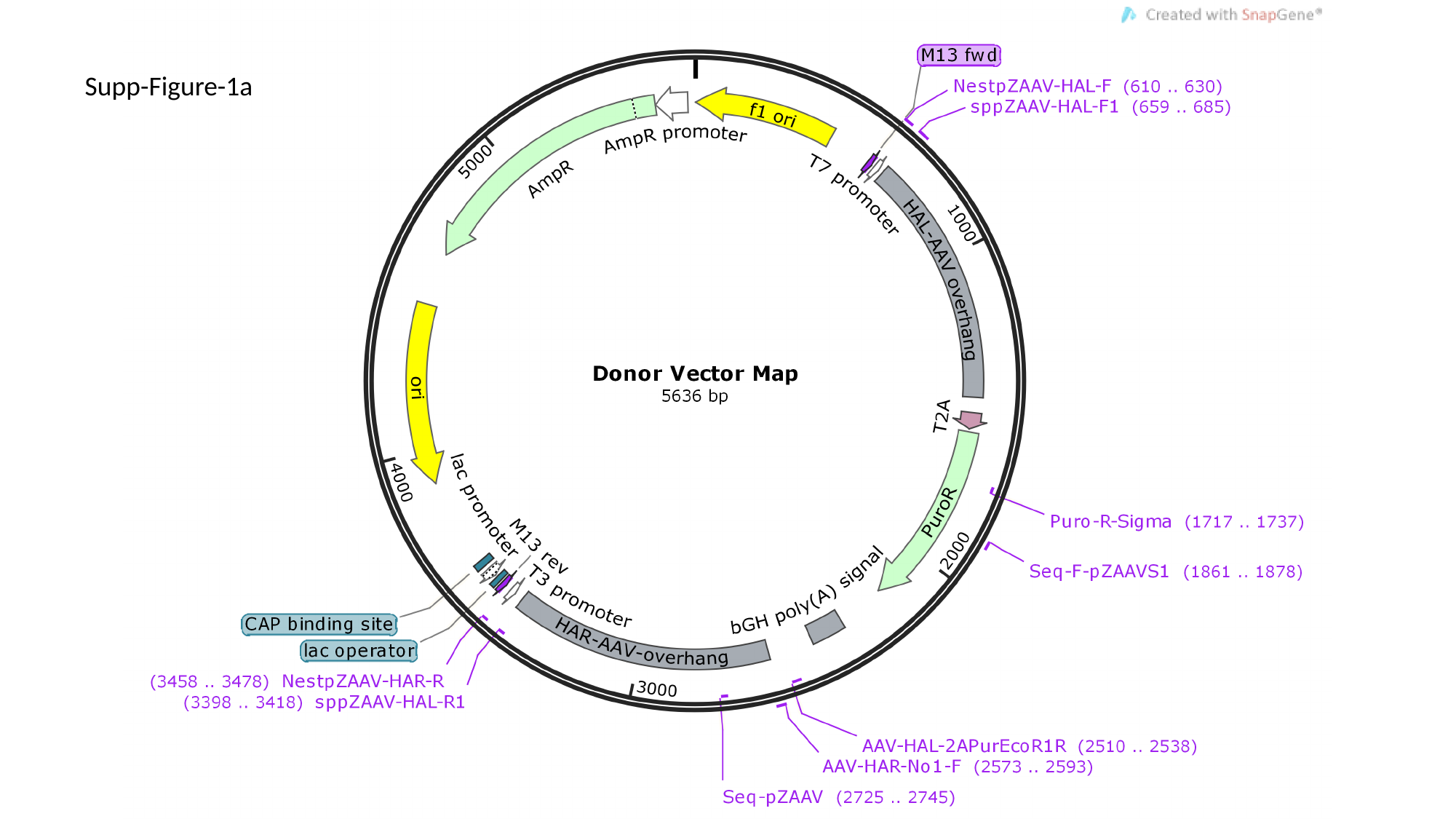

Supp-Figure-1a

### Slide 2
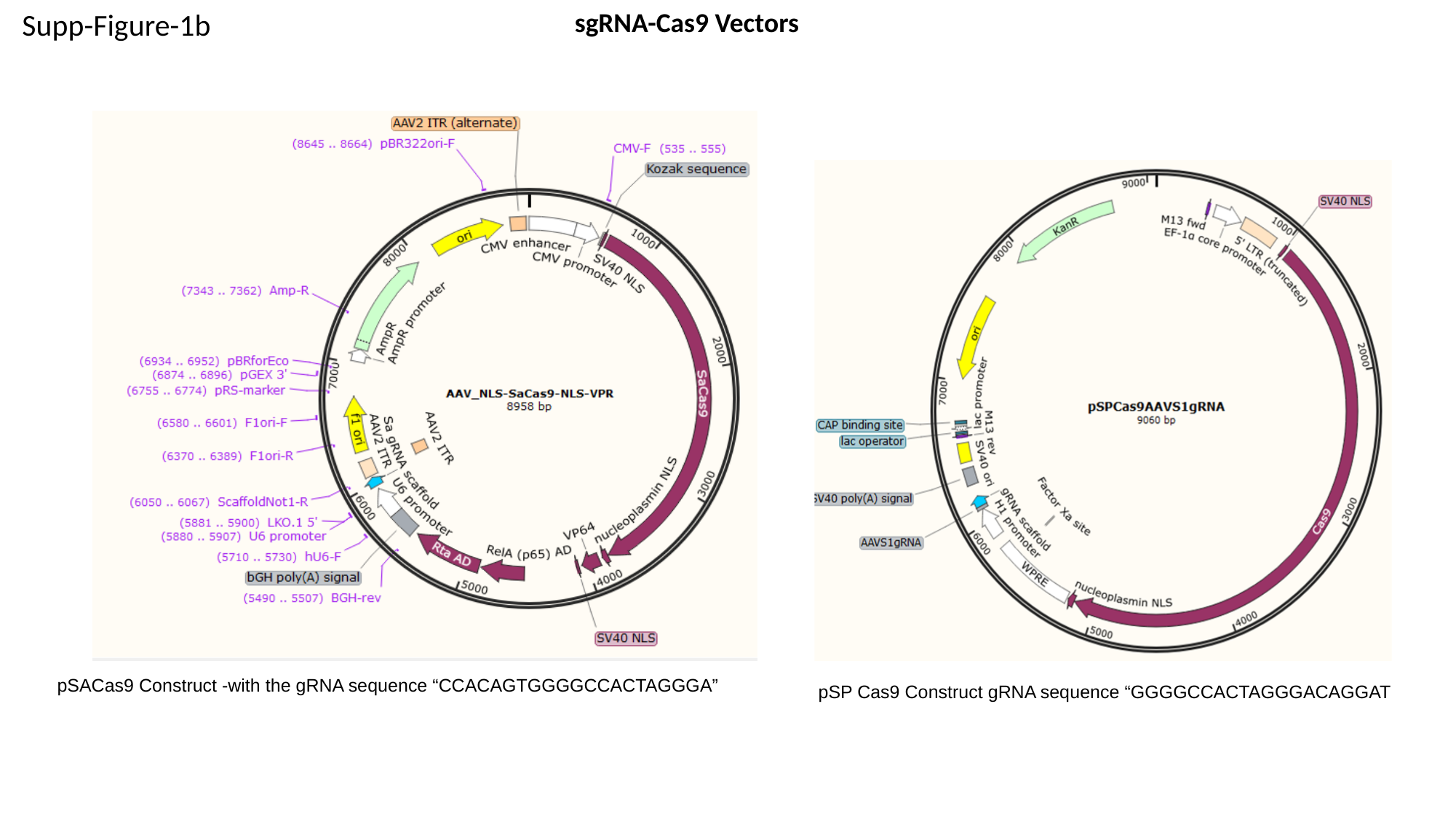

sgRNA-Cas9 Vectors
Supp-Figure-1b
pSP Cas9 Construct gRNA sequence “GGGGCCACTAGGGACAGGAT
pSACas9 Construct -with the gRNA sequence “CCACAGTGGGGCCACTAGGGA”
