## Supplementary material for "Cloning of Human ABCB11 Gene in *E. coli* required the removal of an Intragenic Pribnow-Schaller Box before it’s Insertion into Genomic Safe Harbor AAVS1 Site using CRISPR Cas9": Supp. table

| PRIMER NAME | PRIMER SEQUENCE | Expected Size |
| --- | --- | --- |
| ABCB11 F1 | **ATGTCTGACTCAGTAATTCTTCGAAGTATAAAGAAA** | 1540 bp |
| ABCB11 R1 | **CTGCAATGGTGGTAGAGAACAGAAC** |  |
| ABCB11 F2 | **ATCAGATTGGGATAGTGGAGCAAGAG** | 1597bp |
| ABCB11 R2 | **CACTCAGTACAACTGCAGAGATCAC** |  |
| ABCB11 F3 | **CTGCTTCCTACAGATATGGAGGTTAC** | 997 bp |
| ABCB11 R3 | **TCAACTGATGGGGGATCCAGTG** |  |

Supp.Table-1 ABCB11 PCR amplification primers

Supp. Table-2 ABCB11 sequence we performed a bioinformatic analysis using BPROM to predict bacterial promoters and transcription start-sites

| Promotor position | Threshold for promotor | Number of predicted promotors |  |  |  |  |  |  |  |
| --- | --- | --- | --- | --- | --- | --- | --- | --- | --- |
|  | | Promotor position | LDF | -10 BOX AT POS. | -10 BOX  LENGTH OF SEQUENCE | -10 BOX score | -35 BOX AT POSITON | -35 BOX  LENGTH OF SEQUENCE | -10 BOX score |
|  |  | 91 | 3.84 | 76 | tcatataat | 68 | 53 | atgatg | 22 |
|  |  | 484 | 3.70 | 469 | ggatatatt | 59 | 446 | ttgctg | 47 |
|  |  | 1300 | 2.86 | 1300 | cattatcct | 47 | 1262 | ttgaat | 55 |
|  |  | 983 | 0.87 | 983 | tgggattct | 34 | 951 | tagaaa | 23 |

| PROMOTOR POSITON | TF Binding sites | sequence | Position | score |
| --- | --- | --- | --- | --- |
| 91 | RpoD17 | ATAATAAT | 80 | 8 |
|  | FUR | ATAATGAT | 83 | 6 |
| 484 | lexA | TATATTCA | 472 | 8 |
| 1300 | CRP | TAATGTGA | 1272 | 9 |
| 983 | hns | AAAGGAAT | 955 | 8 |

Supp.Table-3a List of off targets for GGGGCCACTAGGGACAGGAT (PAM TGG)

| Sequence | PAM | Score | #MM | Gene | Locus |
| --- | --- | --- | --- | --- | --- |
| GGGGCCACTAGGGACAGGAT | TGG | N/A |  |  | chr19:-55115751 |
| GGAGACATTAGGGACAGGAT | AAG | 5 | 3 |  | chr10:+119439168 |
| GGGACCATCAGGGACAGGAT | GGG | 11 | 3 |  | chr6:+36797686 |
| GAGGGCAGCAGGGACAGGAT | GGG | 11 | 4 |  | chr12:+131827311 |
| GAGGACAGTAGGGACAGGTT | AAG | 15 | 4 |  | chr18:+8749293 |
| GGGTGCAGTGGGGACAGGAT | GGG | 15 | 4 |  | chr20:-57806974 |
| TGGGCCATCAGGGACAAGAT | GAG | 16 | 4 |  | chr9:+78930561 |
| GAGGGCTCTAGGGACAGGAT | GAG | 18 | 3 |  | chr9:-90086410 |
| AGGG-CACTGGGGACAGGAT | CAG | 18 | 3 |  | chr14:+92970591 |
| GGGGTCACTGGGGACAAGAT | TGG | 20 | 3 |  | chr15:-45535697 |
| GTGGCCAGCTGGGACAGGAT | AGG | 20 | 4 |  | chr9:+34350222 |
| GTGGCCT-TAGGGACAGGAT | AAG | 23 | 3 |  | chr17:-47810510 |
| GGGGTCTTTAGGGACAAGAT | CAG | 23 | 4 |  | chr6:-16277321 |
| GGGGACAG-AGGGACAGGAT | GGG | 25 | 3 |  | chr2:-1587914 |
| GAGGCCA-TAGGGGCAGGAT | GGG | 25 | 3 |  | chr18:-33816007 |
| GGGGACAG-AGGGACAGGAT | GGG | 25 | 3 |  | chr2:-1587705 |
| GTGTCCACCAGGGACAGGTT | GAG | 26 | 4 |  | chr3:-44057223 |
| GGGCACAGTAGGGACAGGAA | GAG | 27 | 4 |  | chr8:-17472019 |
| GGGGCCAATTAGGACAGGAT | GGG | 28 | 3 |  | chr13:+105960562 |
| GGAACCTACTAGGGACAGGAT | GAG | 29 | 3 |  | chr6:-157966897 |
| ATGGCCACTAAGGACAGGAA | AGG | 29 | 4 |  | chr12:+107092486 |
| GGGGTCAGCAGGGGCAGGAT | CGG | 31 | 4 |  | chr17:-75895457 |
| TGGGCCATTAGGGACAAGAA | GGG | 32 | 4 |  | chr15:+63926588 |
| GGGGACA-GAGGGACAGGAT | GGG | 33 | 3 |  | chr19:+45929459 |
| GCAGCCAGGAGGGACAGGAT | GGG | 33 | 4 |  | chr12:+49891213 |
| AGGGTCACTTGAGACAGGAT | CAG | 34 | 4 |  | chr12:-367841 |
| GAGGTCACTGGGGGCAGGAT | AAG | 35 | 4 |  | chr1:-208558678 |
| AGGGGCAGAAGGGACAGGAT | GGG | 35 | 4 |  | chr2:+174691584 |
| GGTGACAGTTGGGACAGGAT | GAG | 36 | 4 |  | chr9:-68603540 |
| GGGGGCACTGAGGGCAGGAT | AAG | 37 | 4 |  | chr21:+21147982 |
| GGTTCCAGCAGGGACAGGAT | CAG | 38 | 4 | BTNL8 | chr5:+180947665 |

Table-3b List of off targets for CCACAGTGGGGCCACTAGGGA (PAM: NNGRRT)

| Sequence | PAM | Score | #MM | Gene | Locus |
| --- | --- | --- | --- | --- | --- |
| CACAGTGGGGCCACTAGGGA | CAG | N/A |  |  | chr19:-55115757 |
| CAC--TGGGGCCACTAGGGA | CAG | 5 | 2 |  | chr5:+96387863 |
| TACAGTGCGGCCACTAAGGA | AGG | 7 | 3 |  | chr6:+43394490 |
| CACAATGGGGACACTAAGGA | GGG | 20 | 3 |  | chr12:+56770752 |
| GACAGTGAGGACCACTAGGGA | GGG | 24 | 3 |  | chr20:+62514542 |
| CAGAGCTGGGCCACTAAGGA | CAG | 25 | 4 | LOC105372863 | chr22:-20232502 |
| CGCA--GGGGCCACTAGGGA | GGG | 27 | 3 |  | chr17:+19111097 |
| CGCA--GGGGCCACTAGGGA | GGG | 27 | 3 |  | chr17:+19237260 |
| AACAGCTGGGTCCACTAGGGA | AAG | 33 | 3 |  | chr13:+106680268 |
| CAGAGTAGGGCCACTATGGA | TAG | 36 | 3 |  | chr2:+76556325 |
| AACAGTGGGGCC-CTAAGGA | AGG | 38 | 3 |  | chr2:-110799006 |
| TAAAGTGGGGCCACTAGGGT | GGG | 38 | 3 |  | chr3:-137074238 |
| CACTCTGGAGCCACTAGGGA | AGG | 38 | 3 |  | chr2:-118261457 |
| AACAGTAGGGCCTCTAGGGA | CGG | 39 | 3 |  | chr7:-64035194 |
| CTCAGT-GGGCCACTAGGTA | CAG | 41 | 3 |  | chr10:+6412824 |
| AACACTGGGGCCAC-AGGGA | CAG | 41 | 3 |  | chr1:-4020013 |
| GACATTGGGACCACTAGAGA | GGG | 42 | 4 |  | chr20:-14875646 |
| CACTGTGGGGCCA-TAGGGA | CAG | 43 | 2 |  | chr10:-112407989 |
| CACAGCAAGGGCCACTAGGGA | AAG | 43 | 3 |  | chr6:+140049644 |
| CTCAGTGTGGTCACTAGGGA | AGG | 44 | 3 | LOC105374577 | chr2:-45331250 |
| AACAATGGGGCCAC-AGGGA | AAG | 44 | 3 |  | chr14:-22144362 |
| CACAGTGAGGCCACTAAGGG | AGG | 44 | 3 |  | chr16:+25422080 |
| CAGAGAGCGCCCACTAGGGA | AAG | 44 | 4 |  | chr3:+124513439 |
| CATATTGGGGCCTCTAAGGA | AGG | 45 | 4 |  | chrX:-110400358 |
| CACAGTAGGGAC-CTAGGGA | GGG | 46 | 3 |  | chr19:-19575574 |
| TACCGTGGGGCCACTAGATA | CAG | 46 | 4 |  | chr2:-15242667 |
| CACATTTGGGAAACTAGGGA | AAG | 48 | 4 |  | chr2:+24766922 |
| GCCAGTGGGGCCACTGGGGA | GGG | 49 | 3 | RALY | chr20:+34080392 |
| CACAGTGTGGCCAC-AGGGA | CAG | 51 | 2 | MYLIP | chr6:-16145268 |
| TACAATGAGGCCATTAGGGA | GGG | 51 | 4 |  | chr4:+125710278 |
| CACTGTGAAGACACTAGGGA | TGG | 51 | 4 |  | chrX:+105863157 |
| AACAGTGGGG--ACTAGGGA | TGG | 52 | 3 |  | chr2:-67797115 |
| CAAAGTGGG-CAACTAGGGA | TGG | 52 | 3 |  | chr10:-7914495 |
| GACTGTGGGATCACTAGGGA | GAG | 52 | 4 |  | chr6:-20120838 |
| CACAGTGGGGACACT-GGGA | GAG | 53 | 2 |  | chr1:-247169522 |
| CACAGTGGGGACACT-GGGA | AAG | 53 | 2 |  | chr11:+133916671 |
| TACACTGAGGCCACTGGGGA | AGG | 53 | 4 |  | chr19:+14553492 |
| TGCAGTGGGGCCCCTAAGGA | CAG | 53 | 4 |  | chr7:-37199736 |

Supp.Table-4 Primers used for validation and sequencing of genomic integration at AAVS1 site

| Primer name | Purpose | Sequence |
| --- | --- | --- |
| HAL-FT7 assay | Sequencing, T7 assay | CATCTCTCCTCCCTCACCCA |
| HAR-R T7 assay | Sequencing, T7 assay | AGGGAGTTTTCCACACGGAC |
| F-AAV-RT-out | Puromycin selection-primary PCR | CCCGTTGCCAGTCTCGAT |
| R-Puro-RT | Puromycin selection-primary PCR | GAGACGCCGACGGTGG |
| F-AAV-HAL-seq-out | Puromycin selection-secondary PCR (nested) | TGACCCATCGAGTCCTCCTT |
| R- Puro | Puromycin selection-secondary PCR (nested) | TGAGGAAGAGTTCTTGCAGCT |
| AAVS1 Forward | PCR Puromycin selection-primary PCR | CGGAACTCTGCCCTCTAACG |
